# Genetic Analysis of Flagellar-Mediated Surface Sensing by *Pseudomonas aeruginosa* PA14

**DOI:** 10.1101/2024.12.05.627040

**Authors:** Sherry Kuchma, C.J. Geiger, Shanice Webster, Yu Fu, Robert Montoya, G.A. O’Toole

## Abstract

Surface sensing is a key aspect of the early stage of biofilm formation. For *P. aeruginosa*, the type IV pili (TFP), the TFP alignment complex and PilY1 were shown to play a key role in c-di-GMP signaling upon surface contact. The role of the flagellar machinery in surface sensing is less well understood in *P. aeruginosa*. Here we show, consistent with findings from other groups, that a mutation in the gene encoding the flagellar hook protein (Δ*flgK*) or flagellin (Δ*fliC*) results in a strain that overproduces the Pel exopolysaccharide (EPS) with a concomitant increase in c-di-GMP levels. We use a candidate gene approach and genetic screens, combined with phenotypic assays, to identify key roles for the MotAB and MotCD stators and the FliG protein, a component of the flagellar switch complex, in stimulating the surface-dependent, increased c-di-GMP level noted for these flagellar mutants. These findings are consistent with previous studies showing a role for the stators in surface sensing. We also show that mutations in the genes coding for the diguanylate cyclases SadC and RoeA as well as SadB, a protein involved in early surface colonization, abrogate the increased c-d-GMP-related phenotypes of the Δ*flgK* mutant. Together, these data indicate that bacteria monitor the status of flagellar synthesis and/or function during surface sensing as a means to trigger the biofilm program.

**Importance:** Understanding how the flagellum contributes to surface sensing by *P. aeruginosa* is key to elucidating the mechanisms of biofilm initiation by this important opportunistic pathogen. Here we take advantage of the observation that mutations in the flagellar hook protein or flagellin enhance surface sensing. We exploit this phenotype to identify key players in this signaling pathway, a critical first step in understanding the mechanistic basis of flagellar-mediated surface sensing. Our findings establish a framework for the future study of flagellar-based surface sensing.

## Introduction

Bacterial biofilms were first formally described in the 1930s (1) and since then this ubiquitous mode of sessile bacterial growth has been shown to be important in both medical and industrial settings (2, 3). The first step in the transition from free swimming planktonic cells to biofilm formation is the microbe contacting the surface and relaying this input signal to the cell to initiate the biofilm mode of growth, a process known as “surface sensing” (4–6).

Many bacteria rely on motility appendages, including flagella and type IV pili, to sense and traverse surfaces. These molecular machines have been shown to be necessary for proper biofilm formation and have been implicated in surface sensing (4–9), but the mechanism(s) by which these appendages sense and transmit the surface sensing signal are just beginning to emerge. Several early studies demonstrated that the bacterial flagellum responds to mechanical load, which in turn can serve as a signal of surface engagement. For example, by manipulating the viscosity of the liquid culture or by adding antibodies specific to the flagellum, surface-associated phenotypes were achieved during liquid culture conditions (10–12) indicating that it is the interference in bacterial flagellum function that is the proximal means whereby microbes detect surface engagement.

Bacterial flagella are used to propel the cell body in both liquid and across surfaces (13). A flagellum is composed of a basal-body structure that spans the cellular envelope in bacteria. A hook and flagellar filament extend from the cell body, and upon rotation, propels the cell body (14). This molecular machine uses ion motive force, generated by a gradient of protons or sodium ions across the cytoplasmic membrane, to rotate the flagellar filament (15–17). This conversion of chemical potential to flagellar rotation is achieved by stator units that can dynamically bind and dissociate from the flagellar motor (18, 19). Stators are composed of an inner membrane (IM) pentamer and a central dimer unit that plugs the ion pore when stators are not incorporated in the flagellum. Upon incorporation into the flagellum, the inner dimer binds the peptidoglycan (PG) layer, unplugging the ion channel within the stator unit, which allows for ion flow down the concentration gradient. This chemical energy is harnessed by the stator units in the form of torque that is transferred to the C-ring of the flagellum via electrostatic interactions with FliG (20, 21). It has been demonstrated that when the flagellar motor experiences a mechanical load, it is able to remodel and recruit additional stator units to aid in rotation (19, 22–24), indicating that changes in external load are sensed by the flagellum and stator occupancy is a readout for this signal.

Recently, studies using different polarly flagellated, monotrichous bacteria have revealed striking similarities in the mechanism by which they use their flagellum to sense surfaces. *Vibrio cholerae*, *Caulobacter crescentus* and *Pseudomonas aeruginosa* have all been used as model organisms to study flagellar-mediated surface sensing and biofilm initiation. One similarity among these model systems is that mutating genes required for flagellar biosynthesis results in surface-associated phenotypes such as exopolysaccharide (EPS) over-production (25–31). Furthermore, enhanced EPS production was dependent on an increase in the second messenger c-di-GMP which was often the result of increased level/activity of one or more diguanylate cyclases (DGCs) (25, 28, 29). Finally, the EPS over-producer phenotype exhibited by different flagellar mutants are not universally dependent on stator function (28, 29), best shown by an exceptionally thorough analysis of flagellar mutant-associated EPS phenotypes and their stator requirements in *V. cholerae* (28). In general, flagellar mutants defective in early stages of flagellar biosynthesis, that is, steps that disrupt assembly of the basal body and motor structures, lack a stator requirement for EPS over-production. In contrast, mutants defective in late stages of flagellar assembly, such as those steps predicted to assemble a basal body and motor but lack a flagellar filament, did require stators for these phenotypes. These observations link the stators to flagellar-mediated surface sensing, particularly when the flagellar machine is almost completely assembled.

While there are similarities in flagellar-mediated surface sensing between *V. cholerae*, *C. crescentus*, and *P. aeruginosa* as described above, the flagellar motor of *P. aeruginosa* is notably distinct among these microbes in that it can accommodate two different sets of stators, MotAB and MotCD. Moreover, these two stator sets have distinct roles in surface motility: MotAB has been shown to be necessary for maximum velocity during swimming motility, whereas MotCD has been shown to be absolutely required for swarming motility (32–35). Additionally, the MotCD stator has been shown to be directly involved in surface sensing by binding to the DGC SadC and stimulating c-di-GMP production upon surface contact (36). This interaction is mediated by the c-di-GMP-binding protein FlgZ when bound to c-di-GMP. The FlgZ·c-di-GMP complex is required for the removal of MotCD stator units from the flagellar motor, leading to shutdown of flagellar rotation while stimulating c-di-GMP production (37). These data indicate that the flagellum, stators, and SadC are important for surface sensing, but there remain missing links in how these complexes are coordinated upon surface contact. The dual-stator system of *P. aeruginosa* may offer unique insights into how bacteria optimize motility and signaling in response to surface engagement.

While proteins involved in flagellum-mediated surface sensing by *P. aeruginosa* have been identified, the mechanism whereby c-di-GMP is increased after initial surface contact by the cell remains a mystery. Here we use a combination of genetic screens and candidate gene studies, combined with phenotypic assays, to begin to investigate how *P. aeruginosa* uses its flagellum to sense a surface.

## Results

### Mutations in the *flgK* and *fliC* genes result in a Pel-dependent increase in Congo red binding and wrinkly colony morphology

A recent publication that included members of our team demonstrated that mutating the gene encoding the hook-associated protein FlgK or the gene encoding the flagellin FliC of *P. aeruginosa* PA14 led to an increase in the production of the Pel EPS by those mutant strains (31). A similar observation was previously made by Parsek and colleagues in *P. aeruginosa* PAO1 when characterizing rugose small colony variants (RSCVs) isolated from biofilm-grown populations and Cystic Fibrosis (CF) sputum (30). When plated on Congo Red (CR) agar, the Δ*flgK* and Δ*fliC* mutants showed enhanced binding of the dye Congo red and a wrinkly colony morphology; these phenotypes were dependent on production of the Pel polysaccharide (38–40) (**Figure 1A, top row**).

**Figure 1.**
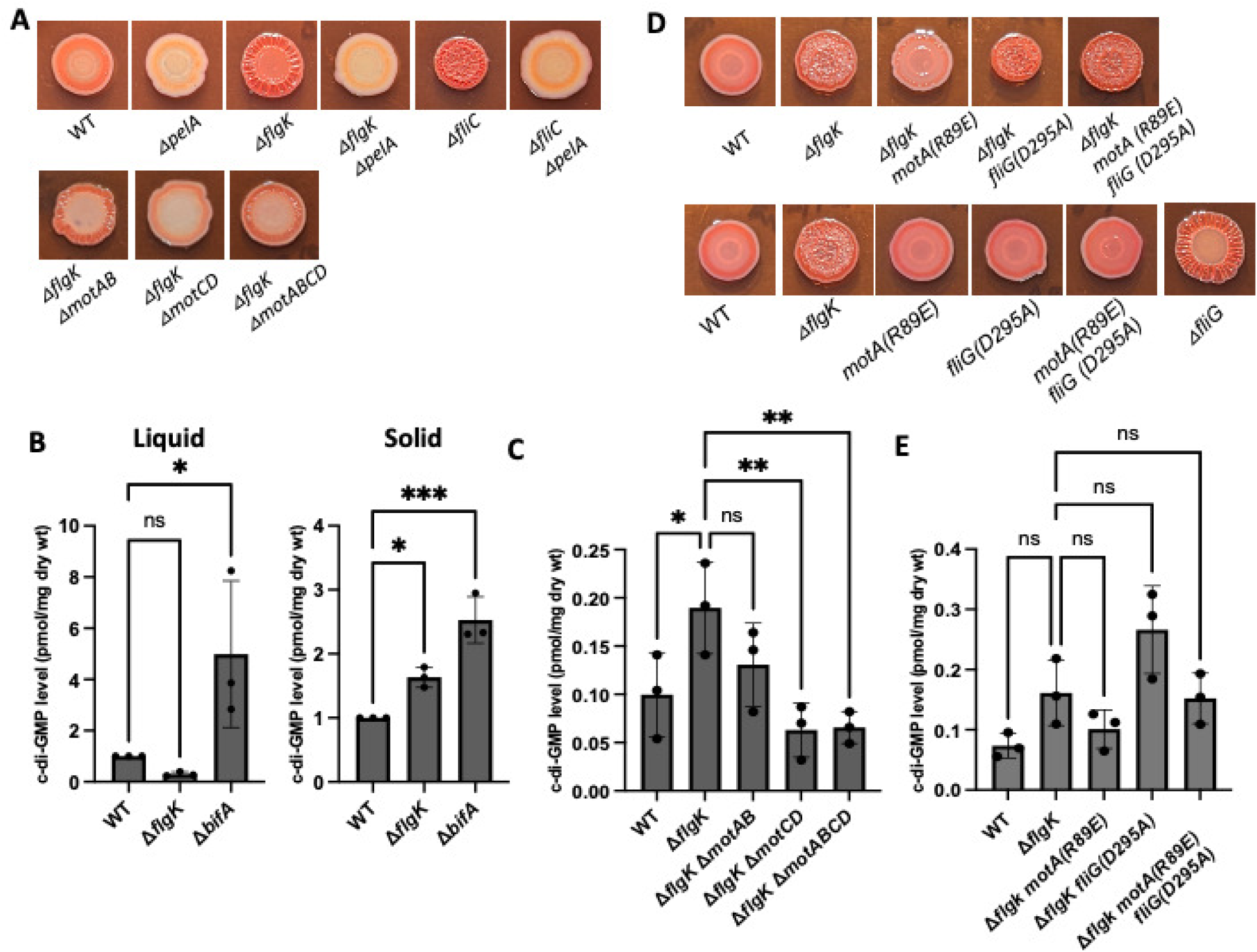
Mutations that eliminate stator production or impact stator occupancy suppress the Δ*flgK* mutant hyper-signaling phenotype. **A and D.** Representative CR images of the indicated strains cultured on M8 medium solidified with 1% agar for 16 h at 37° C followed by 72 hrs at room temperature. **B.** Quantification of c-di-GMP levels in the indicated strains grown either in M8 liquid or on M8 1% agar plates for extraction of nucleotides and measurement of c-di-GMP levels via mass spectrometry. **C and E.** Quantification of c-di-GMP levels in the indicated strains grown on M8 agar swarm plates for 16 h prior to harvest for extraction of nucleotides and measurement of c-di-GMP levels via mass spectrometry. Experiments (**B, C and E**) were performed in triplicate with three technical replicates per strain and analyzed by ANOVA with Dunnett’s multiple comparisons test. Significant differences are shown for comparisons to the Δ*flgK* strain. ns, non-significant difference; * P < 0.05 and ** P < 0.01.

### Identification of factors required for the Congo red phenotypes of the Δ*flgK* mutant using a genetic screen

To gain insight into why the Δ*flgK* mutant exhibited enhanced Pel production, we performed transposon mutagenesis of the Δ*flgK* mutant, then plated the mutants on Congo red medium to evaluate Pel production and colony morphology. We screened approximately 10,000 mutants and identified insertions in 44 genes. **Table 1** shows the mutants that either suppressed or exacerbated the Congo red phenotype of the Δ*flgK* mutant. Many of the transposon insertions mapped to genes required for Pel biosynthesis and secretion machinery, as expected, which served to validate the screen. We also identified mutations mapping to genes required for c-di-GMP production, including the DGC-encoding *roeA* gene, again validating the screen. Finally, the screen identified mutations in genes involved in type IV pili function, with known roles in second messenger signaling (41–45) – we address the implications of these latter findings in the Discussion.

**Table 1.** Mutations identified from Congo red transposon screen in Δ*flgK* mutant background.

| Candidate | Predicted or known function | Congo red phenotype* | Number of alleles isolated |
| --- | --- | --- | --- |
| **Pel biosynthetic operon** |  |  |  |
| *pelB* | forms part of the Pel secretion complex | Reduced | 1 |
| *pelD* | c-di-GMP binding protein; important for Pel secretion | Reduced | 1 |
| *pelA* | Component of the Pel secretion complex | Reduced | 2 |
| *pelE* | Component of the Pel secretion complex | Reduced | 2 |
| *pelC* | Component of the Pel secretion complex | Reduced | 1 |
| *pelF* | Component of the Pel secretion complex | Reduced | 1 |
| *pelG* | Component of the Pel secretion complex | Reduced | 3 |
| **TFP-related functions** |  |  |  |
| *pilQ* | T4P secretin protein | Reduced | 1 |
| *pilW* | Minor pilin, forms the T4P assembly | Reduced | 1 |
| *pilY1* | Important in pili assembly and mechanosensing | Reduced | 7 |
| *pilV* | minor pilin | Reduced | 2 |
| *pilX* | minor pilin | Reduced | 1 |
| **c-di-GMP-related functions** |  |  |  |
| *roeA* | a diguanylate cyclase (DGC) | Reduced | 1 |
| *retS* | Regulator of c-di-GMP level, EPS, T3SS | Enhanced | 1 |
| *bifA-sodB* | Intergenic transposon insertion between *bifA* [phosphodiesterase (PDE)] and *sodB* (superoxide dismutase) | Enhanced | 1 |
| *pvrS* | PvrS part of a two-component system with PvrR (a PDE) | Reduced | 9<sup>#</sup> |

| **Redox-related functions** |  |  |  |
| --- | --- | --- | --- |
| *speA* | arginine decarboxylase | Reduced | 1 |
| *sodM* | superoxide dismutase | Enhanced | 1 |
| *katA* | catalase | Enhanced | 1 |
| PA14_44350 | cbb3-type cytochrome c oxidase subunit II | Enhanced | 1 |
| PA14_57570 | cytochrome c reductase | Enhanced | 3 |
| **Regulators** |  |  |  |
| PA14_16550 | Putative transcriptional regulator | Reduced | 1 |
| PA14_43670 | Histidine kinase, part of a two-component system | Reduced | 1 |
| *hflX* | Role in lysogeny | Reduced | 1 |
| *aguR* | Transcription factor, negative regulation of hydrolase activity | Enhanced | 1 |
| **Other functions** |  |  |  |
| PA14_11290 | Putative permease – membrane transport proteins | Reduced | 1 |
| *thdF* | Putative GTP binding protein - GTPase | Reduced | 1 |
| *ppK* | Polyphosphate kinase – responsible for the synthesis of inorganic polyphosphate from ATP | Reduced | 1 |
| *ptsP* | phosphoenolpyruvate protein phosphotransferase, downstream gene (PA14_04420) has PAS/GGDEF domain, enhanced CR | Enhanced | 1 |
| *hepP* | heparanase | Reduced | 1 |
| PA14_30470 | periplasmic aliphatic sulfonate binding protein | Reduced | 1 |
| PA14_02890 | nucleoside channel forming protein | Reduced | 1 |
| PA14_72870 | aminotransferase, biosynthesis of secondary metabolites | Enhanced | 1 |
| PA14_08600 | 23S rRNA | Enhanced | 1 |
| PA14_08570 | 16S rRNA | Enhanced | 1 |
| *orfN* | NAD-dependent epimerase/dehydrase, glycosylation, group 4 glycosyl transferase | Reduced | 1 |
| PA14_08580 | tRNA-Ile | Enhanced | 1 |
| PA14_40660 | T6SS effector Tse1, amidase activity | Yellow, smooth | 1 |
| PA14_32820 | PA2462 homolog of PAO1 | Enhanced | 1 |
| PA14_70870 | 5s rRNA | Enhanced | 1 |
| PA14_30100 | 50S ribosomal protein L16 3-hydroxylase | Enhanced | 1 |
| PA14_66100 | O-antigen ligase, WaaL, critical for cell wall integrity and motility | Enhanced | 1 |
| *purM* | phosphoribosylaminoimidazole | Enhanced | 1 |
| PA14_41280 | beta-lactamase | Enhanced | 1 |

\* Note – Reduced CR phenotype indicates reduction in both red color binding and wrinkled colony morphology, unless otherwise indicated.

### The increase in c-di-GMP levels for the Δ*flgK* mutant requires surface growth

The increased Congo red binding and wrinkly colony morphology has been associated with increased c-di-GMP levels for *P. aeruginosa* strains and in other organisms with mutations in their flagellar machinery. To assess whether the Δ*flgK* mutant accumulated increased c-di-GMP when grown specifically on a surface, we quantified c-di-GMP levels of surface-grown cells using mass spectrometry and observed that the Δ*flgK* mutant showed a significant increase in c-di-GMP levels relative to the WT (**Figure 1B**, **right**). In contrast, the liquid-grown Δ*flgK* mutant showed a non-significant reduction in c-di-GMP levels relative to the WT (**Figure 1B**, **left**). The Δ*bifA* mutant serves as a positive control for a strain that produces high levels of c-di-GMP under all conditions. When mutated, the *bifA* gene, which codes for a c-di-GMP phosphodiesterase, results in a strain with increased c-di-GMP levels regardless of the presence of a surface. These data suggest that the increased levels of c-di-GMP in the Δ*flgK* mutant requires surface growth.

### The stators are required for the Pel-dependent increase in Congo red binding and wrinkly colony morphology of the Δ*flgK* mutant

Previous studies from our group have shown that the stators play a key role in surface sensing and modulating c-d-GMP levels (32, 35, 36, 46). As previously described in other bacteria, a wrinkly colony/EPS over-producer phenotype of mutants defective in later stages of flagellar synthesis was dependent on the presence of functional stator units (28–30). Given that the flagellar hook-associated protein FlgK is required for later stages of flagellar biosynthesis and Δ*flgK* mutants are expected to assemble a basal body structure, we predicted that enhanced signaling in this mutant would require functional stators. To test this hypothesis, we mutated one or both stator sets in the Δ*flgK* mutant background. Deletion of either set of stators reduced the amount of CR binding and the wrinkled colony phenotype to a similar degree relative to the Δ*flgK* mutant (**Figure 1A, bottom row)**.

We next asked if the loss of Congo red binding and wrinkly colony morphology for the Δ*flgK* mutant carrying the stator mutants was associated with reduced c-di-GMP levels. As shown in **Figure 1C**, the Δ*flgK* Δ*motCD* mutant exhibited a ∼3-fold reduction in c-di-GMP levels relative to the Δ*flgK* parent, whereas the Δ*flgK* Δ*motAB* mutant showed a modest but non-significant reduction in c-di-GMP levels. Deletion of both stator sets in the Δ*flgK* mutant strain (Δ*flgK* Δ*motAB* Δ*motCD*) showed a significant reduction in c-di-GMP levels comparable to the Δ*flgK* Δ*motCD* strain. Taken together, these data indicate that the stators are required for the enhanced EPS production by the Δ*flgK* mutant and that the MotCD stator set may play a more pronounced role in influencing c-di-GMP levels than the MotAB stator set in the context of the Δ*flgK* mutant, an observation that is consistent with our previous findings (36).

### Mutations in the switch complex impact Congo red binding and wrinkly colony morphology of the Δ*flgK* mutant

The interaction of the cytoplasmic portion of MotA with the FliG protein, a member of the rotor (aka, the switch complex, named for its role in switching the direction of rotation of the motor) is thought to be important for stator incorporation into the flagellar motor. Studies in *E. coli* and *Salmonella* have identified key residues in MotA and FliG that are involved in electrostatic interactions between these proteins, which are thought to be critical sites of contact enabling stator incorporation (20, 21, 47). For example, an R90E charge reversal mutation in MotA of *E. coli* led to loss of motility similar to a *motA* deletion, and furthermore, this motility defect was partially rescued by amino acid substitutions that reversed or neutralized the charge of the FliG-D289 residue (21).

Based on the findings in *E. coli*, we introduced the analogous *motA* mutation (R89E) onto the chromosome of *P. aeruginosa* in the native *motA* locus of the WT and the Δ*flgK* strains to ask whether MotA-FliG interactions are required for enhanced Pel production in the Δ*flgK* mutant. We found that the MotA-R89E mutant protein markedly reduced CR binding and colony wrinkling of the Δ*flgK* mutant (**Figure 1D**, **top row**), with no obvious impact on these phenotypes in the WT background (**Figure 1D**, **bottom row**). The R89E mutation also led to a decrease in c-di-GMP levels in the Δ*flgK* strain but this effect was not significantly different when controlling for multiple comparisons (**Figure 1E**). Notably, the R89E mutation has no detectable impact on MotA protein levels, as previously shown (35), discounting the possibility that reduced protein stability impacts these phenotypes.

Next, we introduced a mutation in *fliG* (FliG-D295A) into the WT and Δ*flgK* mutant strains. The D295A mutation is analogous to the D289A substitution of *E. coli,* which rescued the motility defect of the strain expressing the MotA(R90E) mutant protein. We selected the D295A negative charge to neutral substitution to reduce the likelihood that this mutation would also impact electrostatic interactions between FliG and the MotCD stator, which could complicate the analysis. Interestingly, the strain with FliG-D295A mutant protein led to increased CR binding and wrinkling compared to the Δ*flgK* background alone, with no impact on these phenotypes in the WT strain (**Figure 1D-E**). The *fliG* deletion strain (Figure 1D, bottom row, right) which exhibits hyper CR binding (a phenotype we revisit below) is included as a control, confirming that the *fliG*(D295A) mutation is distinct from a null mutant. As expected, given the CR phenotype, the Δ*flgK fliG*(D295A) strain exhibited higher c-di-GMP levels relative to the Δ*flgK* mutant alone, but the increase was not statistically significant after controlling for multiple comparisons.

We then generated the *fliG*(D295A) *motA*(R89E) Δ*flgK* triple mutant strain and observed an intermediate phenotype compared to the double mutant strains, which essentially restored the Δ*flgK* single mutant CR binding and wrinkly colony appearance. This mutant also had c-di-GMP levels similar to the Δ*flgK* mutant. As with the single *motA* (R89E) and *fliG* (D295A) mutants, the double *motA* (R98E) *fliG* (D289A) mutant did not exhibit changes in CR binding or colony morphology in the WT background, indicating that the changes in these phenotypes are specific to the Δ*flgK* background (**Figure 1D-E**). Overall, interactions between the stator and switch complex appear to have a modest impact on c-di-GMP levels, in contrast to the magnitude of change observed for deletion of the stators.

### Mutations that prevent proton binding suppress the Congo red binding, wrinkly colony phenotype and increased c-di-GMP levels in the Δ*flgK* background

Stators generate torque via ion flux through the inner membrane channel formed by the MotAB stator complex when bound to the motor. In *E. coli*, the MotB-D32 residue is considered critical for proton binding and flux through the stator channel (48). As such, a D32A mutation renders MotB unable to bind protons (48), which results in loss of motor occupancy (19). Studies in *Salmonella* showed that MotB-D33N mutant stators were able to associate with the motor but exhibited an increased rate of dissociation relative to wild type MotAB stators (20, 49). Together, these studies indicate that proton binding and/or transport is important for stator incorporation and/or stability in the motor, and thus stator function.

To test whether proton binding ability impacts Δ*flgK* signaling, we constructed the analogous mutation to *E. coli* MotB-D32A in the *P. aeruginosa motB* and *motD* genes (resulting in the amino acid changes D30A and D23A, respectively) and introduced these mutations onto the chromosome at their native loci. We first confirmed that these mutations did not negatively impact protein levels by performing Western blots to detect the His_6_ epitope-tagged MotB or MotD WT and mutant variants (**Figure 2A-B**, middle panels). In fact, the MotB-D30A and MotD-D23A proteins are detected at relatively higher level compared to their WT counterparts. We also observed that the MotB-D30A variant protein migrated more slowly than the WT protein in the SDS-polyacrylamide gel (**Figure 2A**). This difference in migration is not due to a mutation in the coding sequence as confirmed by PCR and sequencing of genomic DNA from the *motB* locus in the WT and MotB-D30A strains (see Materials and Methods for details). Notably, altered mobility of MotB-D32 variants in SDS-polyacrylamide gels has also been previously reported in *E. coli* (48).

**Figure 2.**
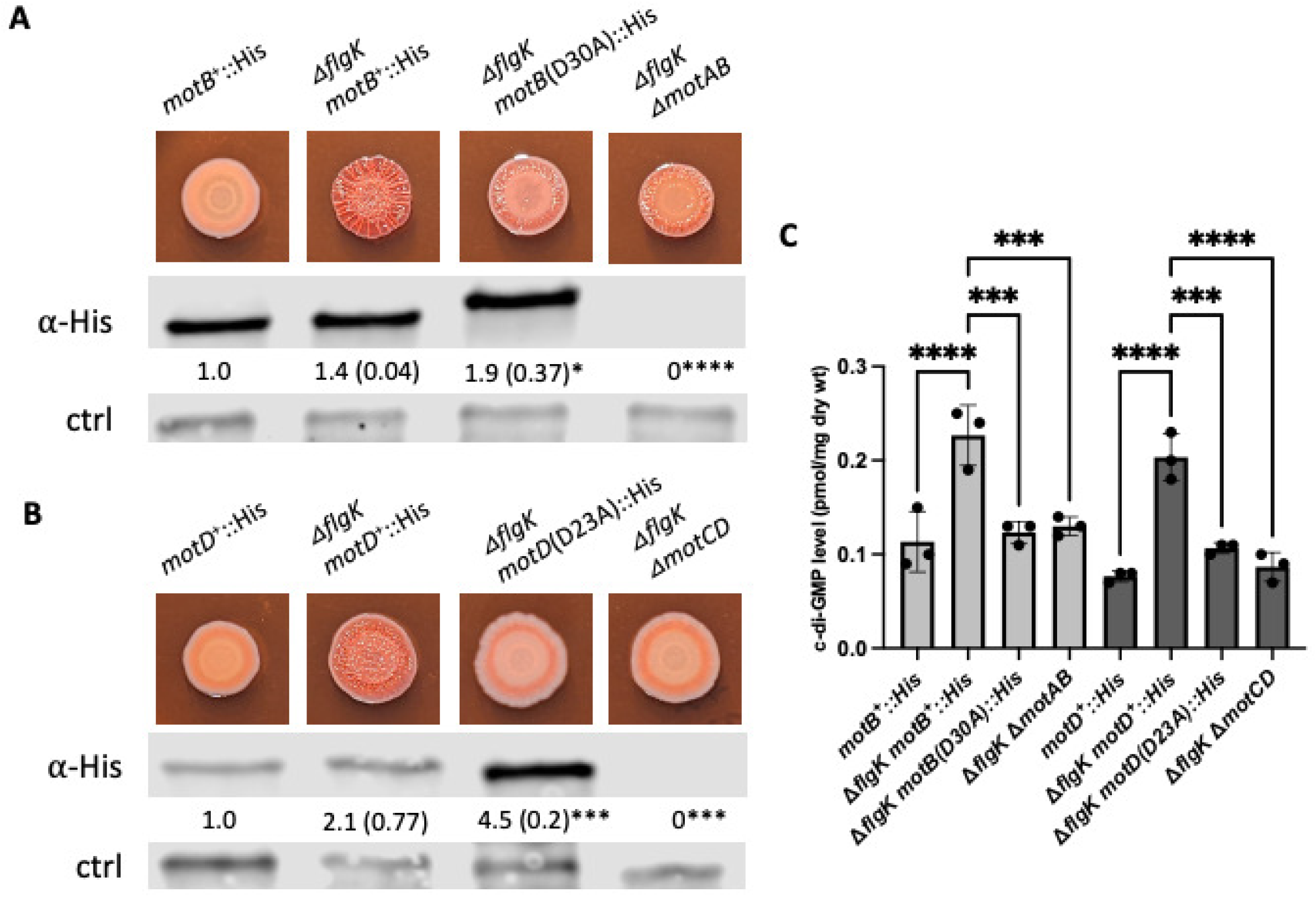
Mutations that impact proton binding suppress hyper-signaling. **A.** Top panel shows representative CR images of the indicated strains. The proton-binding aspartate residue MotB-D30, analogous to the *E. coli* MotB-D32, allele is mutated to alanine in the Δ*flgK* deletion strain. Middle panel shows the Western blot for the MotB-His_6_ WT and D30A variant epitope-tagged proteins detected in lysate samples from surface-grown strains using an anti-His antibody (α-His). MotB-His_6_ protein levels were quantified and normalized to a cross-reacting band (bottom panel, ctrl) detected in all samples and used as a loading control. Numbers below the middle panel show the mean and standard deviation (SD), in parentheses, from three independent experiments, normalized to the WT, which is set to 1.0. Statistical analysis was performed using ANOVA with Dunnett’s test for multiple comparisons. Significant differences are shown for comparisons to the Δ*flgK motB*^+^-His_6_ strain, with * P < 0.05 and **** P < 0.0001. **B.** Top panel shows representative CR images of the indicated strains. The proton-binding aspartate residue of MotD-D23 is mutated to alanine in the Δ*flgK* deletion strain. Middle panel shows the MotD WT and D23A His_6_-epitope tagged proteins detected and quantified as described in panel A. Significant differences are shown for comparisons to the Δ*flgK motD*^+^-His_6_ strain. *, P < 0.05; *** P < 0.0005. **C.** Quantification of c-di-GMP for the indicated strains grown on M8 swarm plates for 16 h. Experiments were performed in triplicate with three technical replicates per strain and analyzed by ANOVA with Tukey’s post-test comparison. Significant differences are shown for comparisons either to the Δ*flgK motB*^+^::His strain or to the Δ*flgK motD*^+^::His strain as indicated. ns, non-significant difference; significant differences noted as follows: ***, P < 0.001; ****, P < 0.0001.

The results showed that both MotB and MotD point mutations required for ion flux phenocopied the deletion mutants of the respective stator sets for both CR binding, wrinkly colony morphology and for c-di-GMP levels (**Figure 2A-C**), indicating that the proton-binding, likely via stator occupancy of MotB and MotD, is important for their role in increased signaling by the Δ*flgK* mutant. Taken together with the switch complex data above, these data are consistent with the previous findings that stator occupancy in the motor is required for signaling.

### The DGCs SadC and RoeA are required for the enhanced c-di-GMP levels in the Δ*flgK* mutant

Previous studies from our team have implicated the SadC and RoeA DGCs as key for early biofilm formation and surface sensing (36, 42, 45, 50–52). Also, as noted above, a transposon mutation in the *roeA* gene was isolated in the Δ*flgK* CR screen as a suppressor of the enhanced CR binding phenotype. Therefore, we asked whether null mutations in the *sadC* or *roeA* genes could impact the Congo Red, wrinkly morphology or c-di-GMP levels of the Δ*flgK* mutant. Mutating the *sadC* or *roeA* genes individually reduced the Congo Red and wrinkly morphology phenotypes of the Δ*flgK* mutant (**Figure 3A**) with the *roeA* mutation having a stronger impact on both phenotypes. Both mutations led to significant reductions in c-di-GMP levels relative to the Δ*flgK* mutant (**Figure 3B**). The triple Δ*flgK* Δ*sadC* Δ*roeA* mutant showed a further reduction in CR binding and wrinkly colony phenotypes compared to each of the double mutants, although the change is modest compared to the Δ*flgK* Δ*roeA* mutant (**Figure 3A-B**). Together, these data indicate that the c-di-GMP produced in the Δ*flgK* mutant is largely contributed by RoeA and SadC.

**Figure 3.**
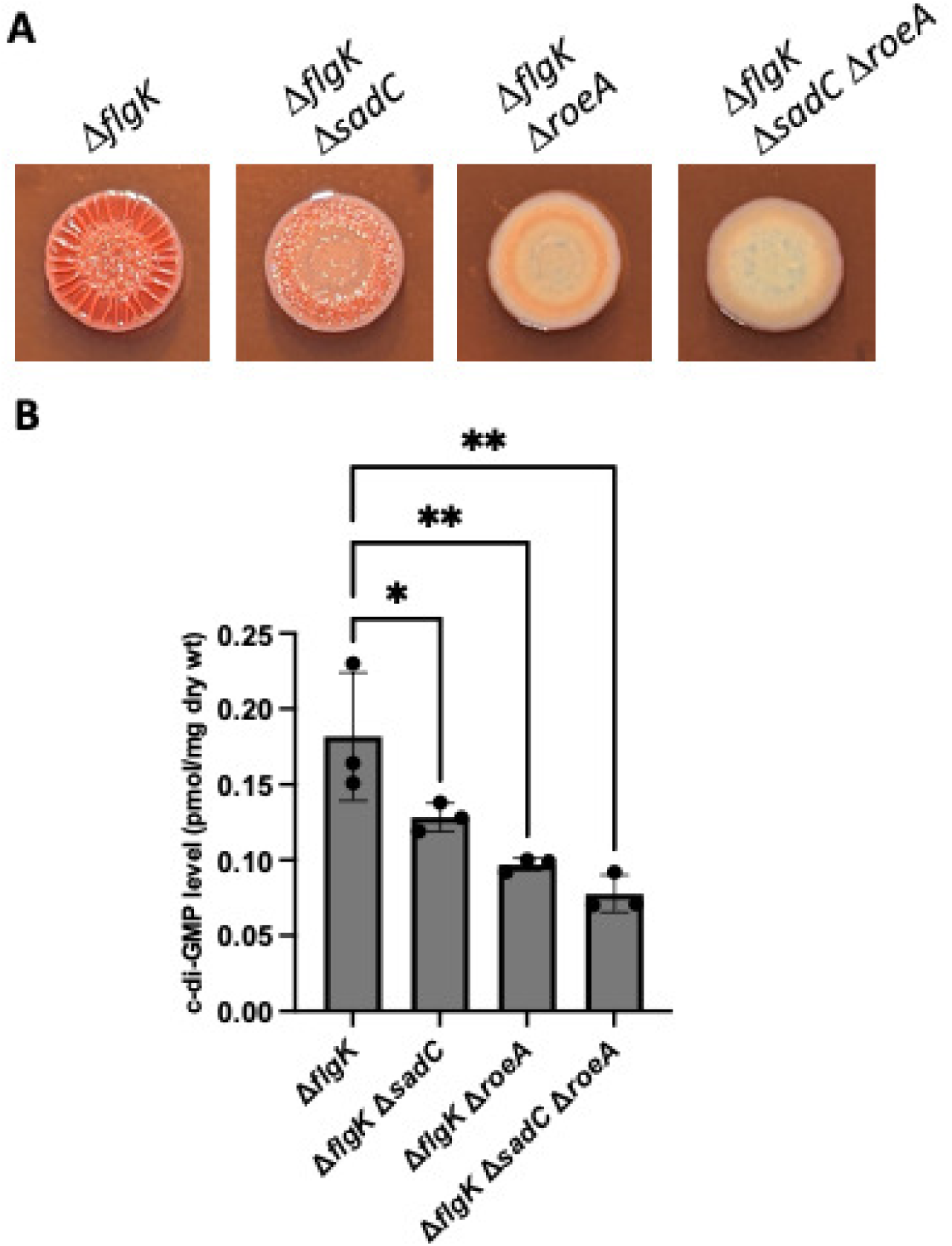
The DGCs SadC and RoeA are required for hyper signaling in the *flgK* mutant. **A.** Representative CR images of the indicated strains. **B.** Quantification of c-di-GMP extracted from swarm-grown strains. Experiments were performed in triplicate with three technical replicates per strain and analyzed by ANOVA with Dunnett’s post-test comparison. Significant differences are shown for comparisons to the Δ*flgK* mutant; *P < 0.05, **P < 0.005.

### Testing candidate genes for their impact on phenotypes of the Δ*flgK* mutant

In addition to the screens described above, we took a candidate gene approach to identify additional genetic factors that may contribute to the Δ*flgK* mutant phenotypes (**Figure 4**). We assessed a number of accessory flagellar factors with known roles in flagellar assembly and function such as FlhF, required for positioning of the flagellum at the cell pole (53–55) and FliL, an accessory protein with a myriad of supporting roles depending on the microbe (16, 18, 56, 57). Mutations in these genes did not alter the Δ*flgK* mutant phenotypes (**Figure 4A**, **top row**), indicating they are not required for enhanced CR binding and wrinkly colony morphology, nor did single mutations in *flhF* or *fliL* genes exert differences in CR binding or colony phenotypes with respect to the WT, indicating these flagellum-related mutations do not impact Pel production (**Figure 4A**, **bottom row**).

**Figure 4.**
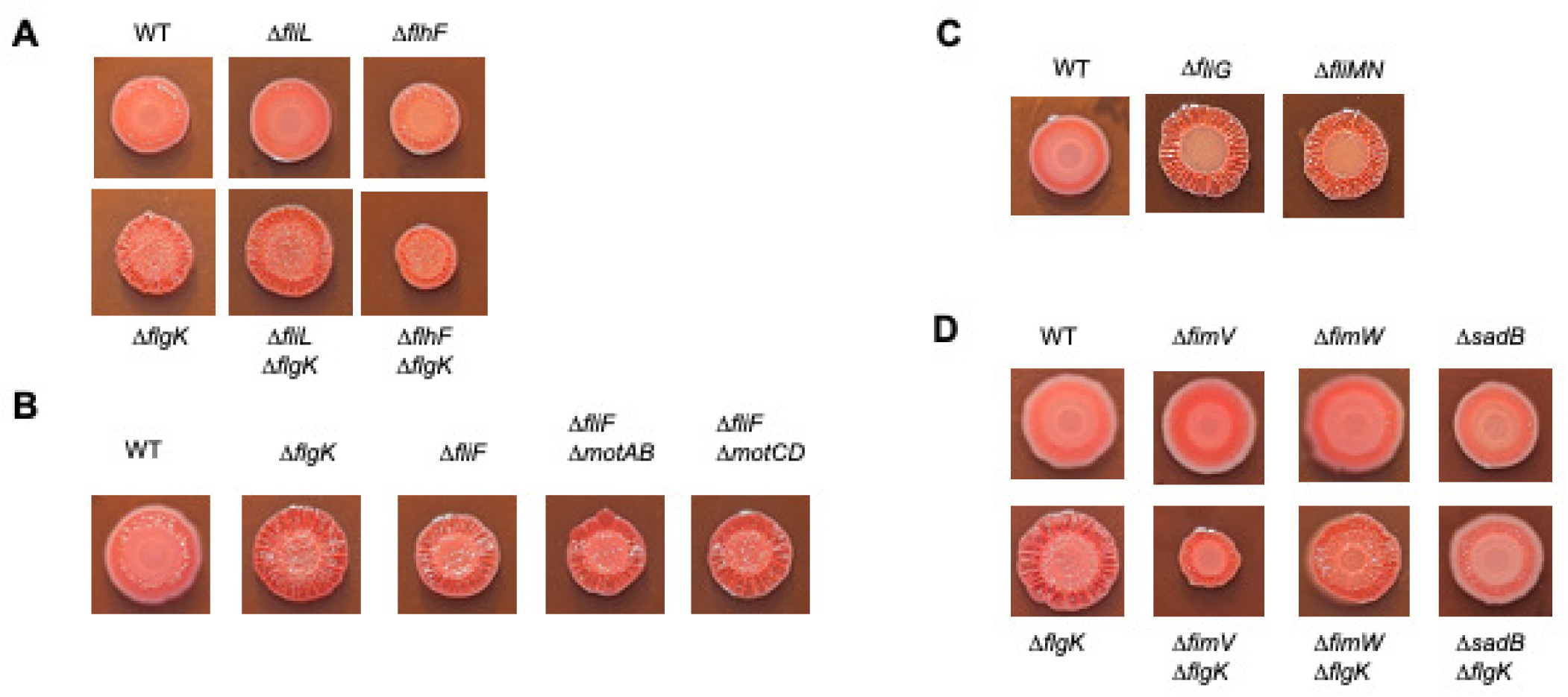
Testing candidate genes for their impact on the Δ*flgK* mutant hyper-signaling phenotype. **A.** Representative CR plate images of the Δ*fliL and* Δ*flhF* mutations in the WT and the Δ*flgK* mutant background. **B.** Impact of mutations in the stators on the Δ*fliF* mutant phenotypes. **C.** Representative CR plate images of mutations the *fliG* and *fliMN* genes. **D.** Impact of mutations in the *fimV*, *fimW*, and *sadB* genes in the WT and Δ*flgK* mutant backgrounds on the CR plate phenotype. All CR assays were performed on M8 medium solidified with 1% agar for 16 h at 37° C followed by 72 hrs at room temperature.

In contrast, mutating the *fliF* gene which encodes the protein comprising the MS (membrane-supra-membrane) ring of the flagellar basal body (56) led to an increase in CR binding and colony wrinkling in the WT background, but the increase is less robust relative to the Δ*flgK* mutant (**Figure 4B**). Mutation of *fliF* is expected to disrupt the basal body structure and preclude motor assembly and stator incorporation. Thus, the Δ*fliF* mutant falls into a class of flagellar mutants that impact early stages of flagellar biosynthesis and have been shown to enhance EPS production in a stator-independent manner in other microbes (28, 29). Based on those findings, we assessed whether the stators were required and found that the Δ*fliF* mutant phenotypes are indeed stator-independent, as mutations in *motAB* or *motCD* did not alter the Δ*fliF* enhanced CR binding and colony wrinkling phenotypes (**Figure 4B**).

We next assessed the impact of deleting the *fliG* and *fliMN* genes in the WT strain background. Mutating the gene coding for FliG, the rotor component that interacts with the stator complex, or FliMN proteins which, together with FliG, make up the switch complex to control the direction of flagellar rotation (58), results in phenotypes similar to the Δ*flgK* mutant (**Figure 4C**). The enhanced CR binding and wrinkled colony phenotypes we observe for the *fliG* and *fliMN* mutant phenotypes are consistent with those previously reported for these mutants in other microbes, and like mutation of the *fliF* gene, these mutations were shown to trigger stator-independent signaling (28, 29).

The CR genetic screen described above (**Table 1**) identified factors required for Δ*flgK* mutant hyper-signaling phenotype that are related to TFP production/function and previously shown to be involved in surface sensing, such as PilY1 (41, 44), PilW and PilX (59). To further assess the roles of additional TFP-related genes with reported roles in c-di-GMP-related signaling, we tested mutations in the *fimV* gene encoding a TFP-associated peptidoglycan binding protein shown to interact with and influence activity of the DgcP diguanylate cyclase (60–63) and the *fimW* gene encoding a c-di-GMP receptor shown to be involved in early cell-surface commitment (64). Our results show that neither mutation of *fimV* nor *fimW* impacted the Δ*flgK* phenotypes (**Figure 4D**), indicating these factors are not required for enhanced signaling in the Δ*flgK* mutant.

Finally, we assessed whether SadB, a protein required for the transition of cells from reversible to irreversible attachment during early stages of surface association, is necessary for the Δ*flgK* mutant phenotypes. Loss of SadB results in enhanced motility, loss of biofilm formation and suppression of the hyper-biofilm and Pel-mediated wrinkly morphology phenotype associated with loss of the BifA phosphodiesterase (65–67). Here we found that the Δ*flgK* Δ*sadB* double mutant reversed the CR-binding and wrinkly phenotypes of the Δ*flgK* mutant (**Figure 4D**), indicating that *sadB* is required for enhanced signaling by the Δ*flgK* mutant. These data agree with previous findings showing that *sadB* mutations suppress the RSCV phenotypes of *fliC* mutants (30). While the precise function of SadB is not yet known, the data here showing that SadB contributes to the flagellum-mediated surface signaling pathway are consistent with its role as an important player in the inverse regulation of motility and biofilm formation (67).

## Discussion

In this study we explored the observation that a mutation in the *flgK* gene results in an increase in c-di-GMP levels as well as an increase in Congo red staining and a wrinkly colony morphology. The observation that the increase in c-di-GMP levels in the Δ*flgK* mutant only occurs on a surface suggested a link to a surface-sensing pathway. To explore the link to surface sensing by leveraging the Δ*flgK* mutant, we performed a genetic screen that identified multiple loci that reduced the Congo red staining and wrinkly colony morphology of this strain.

From the screen, we identified expected pathways (e.g., Pel EPS synthesis) as well as loci that have been linked previously to surface sensing. That is, in a previous study, we performed a genetic screen starting in the *P. aeruginosa* PA14 Δ*bifA* mutant background -the *bifA* gene encodes a c-di-GMP phosphodiesterase (PDE) and the Δ*bifA* mutant produces levels of c-di-GMP ∼10-fold higher than the WT and cannot swarm (51, 65). We mutagenized the Δ*bifA* mutant with the mariner transposon and screened ∼5500 mutants to identify those with restored swarming motility. The list of candidates identified in that previous screen overlapped the candidate mutants identified here, either in the Δ*flgK* transposon screen or the candidate mutants we analyzed, including mutations in genes coding for the stators (*motA*), TFP-related functions linked to cAMP signaling and surface sensing (*pilY1*, *fimU*), c-di-GMP-related functions (PvrRS, a PDE/DGC pair with alleles frequently isolated as suppressors due to Mariner-based over-expression of PvrR as we previously reported), *sadB* and Pel biosynthesis (*pelA*, *pelB*, *pelF*). This finding is perhaps not surprising given that both the Δ*bifA* and Δ*flgK* mutants have increased c-di-GMP levels. These data are consistent with the findings here that mutations in the same genes which reduce the Congo red staining and wrinkly colony morphology of the Δ*flgK* mutant also restored swarming motility to the Δ*bifA* mutant, and previous observations showing the reciprocal regulation of biofilm-versus motility-related functions (50, 67).

We and others have shown previously that T4P are key players in surface sensing (41–45, 68, 69). We note that multiple mutations impacting the T4P were identified as suppressors of the enhanced CR binding and wrinkly colony c-di-GMP-mediated phenotypes of the Δ*flgK* mutant in the CR screen described above. We take this finding to mean that inputs from both flagella and T4P are needed to fully engage the surface sensing pathway. This conclusion is also consistent with our observation that the Δ*flgK* mutant requires surface engagement for increased levels of c-di-GMP. Interestingly, certain mutations that impact T4P, namely *fimV* and *fimW* examined above, do not suppress the Δ*flgK* hyper signaling phenotypes, hinting at the complexity by which the flagellar-and T4P-mediated signals may be integrated to coordinate surface sensing which remains an open question.

We also showed that the DGCs RoeA and SadC contribute to elevated c-di-GMP levels and enhanced signaling phenotypes of the Δ*flgK* mutant. RoeA and SadC have both been shown previously to contribute to biofilm formation, and SadC is a key component of the surface sensing pathway in *P. aeruginosa* PA14 (36, 42, 45, 51, 52). Here, we observed that mutation of the *roeA* gene exerted a stronger impact on Pel-mediated phenotypes of the Δ*flgK* mutant versus Δ*sadC* mutation, although they both showed significant reductions in c-di-GMP levels. This difference in strength of suppression may be one reason we isolated an allele of *roeA* but not *sadC* in the Δ*flgK* CR suppressor screen. Additionally, the absence of *sadC* suppressor alleles (as well as other expected alleles such as those of *sadB*) may be due to the screen not reaching full saturation. Such limitations served as the rationale for targeting candidate genes as a companion approach. These results also agree well with data from a previous report from our group pertaining to suppression of the Δ*bifA* mutant phenotypes, with RoeA playing a stronger role in impacting increased EPS production whereas SadC had a more robust impact on motility repression by the Δ*bifA* PDE mutant (51). Furthermore, those observations dovetail with our findings that SadC interacts with the stator MotC (36); this interaction serves to stimulate SadC’s production of c-di-GMP and thus likely contributes to flagellar signaling. There are now numerous reports of DGCs and PDEs that directly interact with signaling effectors/partners to achieve localized impacts on signaling outcomes (60, 70–72), highlighting a common theme emerging for c-di-GMP signaling pathways.

Our findings also support a role for the stators and switch complex in the phenotypes associated with the Δ*flgK* mutant. Loss of the stators and mutations that render the stators unable to conduct protons result in loss of Congo red binding and the wrinkly colony phenotype of the Δ*flgK* mutant. Similarly, mutations that impact stator interactions with the switch complex also impact CR binding and wrinkly colony appearance of the Δ*flgK* mutant. Here we tested the stator-switch complex interaction using specific allelic combinations of *motA* and *fliG* designed to first disrupt and then restore key electrostatic interactions between these proteins. The design of these mutations was based on studies in *E. coli*. Our results are largely consistent with the notion that disruption of MotA-FliG interaction by *motA* point mutation leads to abrogation of the Δ*flgK* mutant phenotypes, and restoration of the interaction by *fliG* point mutation restores the Δ*flgK* mutant phenotypes. Unexpectedly, we observed that the *fliG*-D295A allele used in these studies enhanced the hyper signaling phenotype in the Δ*flgK* (*motA*^+^) background, rather than the expected result of having little or no impact on the Δ*flgK* phenotypes in this strain. We interpret these observations to indicate that the FliG-D295A protein enhances interactions with the WT MotA protein leading to increased signaling, however, further work is needed to understand the impact of this *fliG* mutation on the signaling pathway.

Collectively, this work, together with studies from diverse bacterial model systems, clearly show there are conserved signaling pathways connecting the flagellum and flagellum biosynthesis to biofilm-relevant signaling. Additionally, another important conclusion from these studies is the notion that bacteria monitor the status of flagellar synthesis as well as flagellar function, such that disruptions in flagellar biosynthesis trigger biofilm-related signaling. In general, late disruptions in flagellar synthesis (after basal body assembly, as in the case for the *fliC* and *flgK* mutants presented here and elsewhere), result in enhanced stator-dependent c-di-GMP signaling and we believe such effects functionally mimic the surface sensing process with cells interpreting lack of the flagellar filament as an increased load on the motor upon surface engagement. In contrast, early disruption in flagellum biosynthesis that precludes basal body production generally leads to enhanced c-di-GMP signaling in a stator-independent manner (as in the case of *fliF* mutation), indicating that bacteria utilize c-di-GMP-mediated signaling to respond to aberrant or aborted flagellar synthesis and thereby promote biofilm formation under such circumstances. Together, these studies highlight the importance of understanding the role of the flagellum in surface sensing and biofilm formation and call for additional studies aimed at further exploration of this salient topic. Key questions remain regarding various aspects of the surface sensing process, including: how does stator function influence c-di-GMP production at the molecular level? What are the mechanistic features that distinguish the MotAB and MotCD stator sets and their roles in transducing the flagellar surface sensing signal? What additional regulatory proteins or pathways might modulate stator-mediated signaling? These and related questions are the focus of current and future work from our group.

## Materials and Methods

### Strains and media

*P. aeruginosa* UCBPP PA14 was used as the WT strain and all mutations were made in this background unless stated otherwise. Mutations were made using *E. coli* S17-1 λpir. All strains used in this study are listed in **Table S1**. Bacterial strains were cultured in 5 ml of lysogeny broth (LB) medium or plated on 1.5% LB agar with antibiotics, when necessary. Gentamicin (Gm) was used at 25 or 30 µg/ml for *P. aeruginosa* and 10 µg/ml for *E. coli*. Carbenicillin (Cb) was used at 100 ug/ml for *E. coli* and Triclosan was used at 20 ug/ml for counter-selection against *E. coli* after conjugation with *P. aeruginosa*. M8 minimal salts medium supplemented with MgSO_4_ (1mM), glucose (0.2%) and casamino acids (0.5%) was used for all assay conditions (73).

### Construction of mutant strains and plasmids

Plasmids used in this study are listed in **Table S2** and primers used in this study are listed in **Table S3**. Plasmids to generate gene deletions and point mutations were generated by PCR and Gibson assembly (74) and cloned into the pMQ30 vector (75). In-frame deletions and point mutants were generated using allelic exchange as previously described (75, 76).

### Transposon mutagenesis and identification of integration site

Transposon mutants were generated with the Mariner transposon as previously described (77). Briefly, *E. coli* S17 harboring the pBT20 plasmid harboring the Mariner transposon was co-incubated with *P. aeruginosa* PA14 Δ*flgK* on LB agar for 1 hour at 30°C for conjugation to occur. Cells were then scraped-up, diluted, and plated on LB agar plates supplemented with 30 µg/ml Gm, 20 µg/ml Triclosan, 0.04 mg/ml Congo Red, and 0.01 mg/ml Coomassie blue. Plates were then incubated at 37C for 24 hours and then at room temperature for 48 hours. Colonies that displayed altered colony morphology or Congo Red uptake relative to the Δ*flgK* strain were selected and confirmed with a second round of plating on Congo Red agar with selection. After confirmation of the phenotype, arbitrary primed PCR was then performed and sequenced using the Sanger method to identify the location and direction of the transposon as previously described (7).

### Congo Red assay

Congo red dye uptake was adapted from previously published protocols (38). Briefly, M8 agar (1 %) plates supplemented with Congo Red solution (final concentration CR at 0.04 mg/ml with 0.01 mg/ml Coomassie blue) were spotted with 2 ul of an overnight culture and incubated at 37°C for 16 hours and then at room temperature for 72 to 96 hours.

### Swarming motility assay

Swarming assays were performed as previously described (78). M8 medium was supplemented with 0.5 % agar. Swarm plates were inoculated with 2 ul of an overnight culture and incubated at 37° C for 16 hours.

### Protein detection and quantification

Cells were harvested from swarm plates grown for 16h at 37°C, normalized by OD_600_ and lysed by boiling in SDS gel loading buffer (50 mM Tris-HCl pH 6.8, 2% SDS, 10% glycerol, 0.1 % bromophenol blue, and supplemented with freshly added DTT at a final concentration of 100 mM). Equal volumes of sample lysates were then resolved using 10 % SDS-PAGE TGX gels (Bio-Rad, Hercules, CA). Proteins were transferred to a nitrocellulose membrane using a Trans-Blot Turbo system (Bio-Rad, Hercules, CA) and probed with anti-His antiserum (Qiagen, Germantown, MD) at 1:2000 dilution prepared in 1X TBS, 3 % BSA. Proteins were detected using fluorescence detection with IRDye-labeled fluorescent secondary antibodies and imaged using the Odyssey CLx Imager (LICOR Biosciences, INC., Lincoln, NE). Image Studio Lite software (LICOR Biosciences, Inc., Lincoln, NE) was used to quantify protein levels using a non-specific band present in all lanes as a normalization control. During Western blotting of the MotB WT and D30A variant, we noticed a change in mobility of the D30A variant protein relative to the WT protein. This mobility shift was observed in both the *flgK* mutant (shown in Figure 2A) and in the WT background (data not shown). One possible explanation for a size shift is a change in the MotB DNA coding sequence, such as a DNA insertion or a mutation of the stop codon leading to read-through into downstream sequence. We investigated these possibilities by sequencing a PCR product amplified from the genomic region encompassing the *motB* gene as well as upstream and downstream sequences in both the WT and variant *motB* genes in the WT and Δ*flgK* strains, however, we did not find any mutations consistent with this explanation. An alternative possibility that has yet to be explored is that the MotB-D30A protein undergoes post-translational modification that alters its gel migration.

### Cyclic-di-GMP quantification

c-di-GMP levels were quantified via liquid chromatography-mass spectrometry (LC-MS) at the Michigan State University Mass Spectrometry and Metabolomics Core. For surface-grown c-di-GMP measurements, cells were harvested from either swarm plates after ∼16 hrs of growth or after 5h growth on 1% agar M8 plates (where noted here) as previously reported (44, 59). For liquid-grown c-di-GMP measurements, cells were sub-cultured (1:100) from an LB-grown overnight culture in M8 liquid media and harvested after 8 hrs at 37°C. Measurements were normalized to dry weight of cell pellets after nucleotide extraction. All experiments were performed in triplicate with three technical replicates per strain.

### Statistical methods

Data were analyzed by ordinary one-way ANOVA with a post hoc test for multiple comparisons (the specific test used for each case is identified in the figure legend) using Graphpad Prism software (La Jolla, CA).

## Acknowledgements

This work was supported by NIH grants R37/AI83256 to GAO and R01/AI143730 to GCLW.

## Literature Cited

1. Henrici AT. 1933. Studies of freshwater bacteria: I. A direct microscopic technique. J Bacteriol 25:277–287.

2. Sharma S, Mohler J, Mahajan SD, Schwartz SA, Bruggemann L, Aalinkeel R. 2023. Microbial biofilm: a review on formation, infection, antibiotic resistance, control measures, and innovative treatment. Microorganisms 11:1614.

3. Zobell CE, Allen EC. 1935. The significance of marine bacteria in the fouling of submerged surfaces. J Bacteriol 29:239–251.

4. Belas R. 2014. Biofilms, flagella, and mechanosensing of surfaces by bacteria. Trends Microbiol 22:517–527.

5. Chawla R, Gupta R, Lele TP, Lele PP. 2020. A skeptic’s guide to bacterial mechanosensing. J Mol Biol 432:523–533.

6. O’Toole GA, Wong GC. 2016. Sensational biofilms: surface sensing in bacteria. Curr Opin Microbiol 30:139–146.

7. O’Toole GA, Kolter R. 1998. Flagellar and twitching motility are necessary for *Pseudomonas aeruginosa* biofilm development. Mol Microbiol 30:295–304.

8. Ellison CK, Kan J, Dillard RS, Kysela DT, Ducret A, Berne C, Hampton CM, Ke Z, Wright ER, Biais N, Dalia AB, Brun YV. 2017. Obstruction of pilus retraction stimulates bacterial surface sensing. Science 358:535–538.

9. Ellison C, Brun YV. 2015. Mechanosensing: A regulation sensation. Curr Biol 25:R113– R115.

10. Cairns LS, Marlow VL, Bissett E, Ostrowski A, Stanley-Wall NR. 2013. A mechanical signal transmitted by the flagellum controls signalling in *Bacillus subtilis*. Mol Microbiol 90:6–21.

11. McCarter L, Hilmen M, Silverman M. 1988. Flagellar dynamometer controls swarmer cell differentiation of *V. parahaemolyticus*. Cell 54:345–351.

12. McCarter L, Silverman M. 1990. Surface-induced swarmer cell differentiation of *Vibrio parahaemoiyticus*. Mol Microbiol 4:1057–1062.

13. Jarrell KF, McBride MJ. 2008. The surprisingly diverse ways that prokaryotes move. Nat Rev Microbiol 6:466–476.

14. Aldridge P, Hughes KT. 2002. Regulation of flagellar assembly. Curr Opin Microbiol 5:160– 165.

15. Deme JC, Johnson S, Vickery O, Aron A, Monkhouse H, Griffiths T, James RH, Berks BC, Coulton JW, Stansfeld PJ, Lea SM. 2020. Structures of the stator complex that drives rotation of the bacterial flagellum. Nat Microbiol 5:1553–1564.

16. Guo S, Liu J. 2022. The bacterial flagellar motor: Insights into torque generation, rotational switching, and mechanosensing Front Microbiol 13:911114.

17. Homma M, Kojima S. 2022. The periplasmic domain of the ion-conducting stator of bacterial flagella regulates force generation. Front Microbiol 13:869187.

18. Subramanian S, Kearns DB. 2019. Functional regulators of bacterial flagella. Annu Rev Microbiol 73:225–246.

19. Tipping MJ, Delalez NJ, Lim R, Berry RM, Armitage JP. 2013. Load-dependent assembly of the bacterial flagellar motor. MBio 4:10–1128.

20. Morimoto YV, Nakamura S, Kami-ike N, Namba K, Minamino T. 2010. Charged residues in the cytoplasmic loop of MotA are required for stator assembly into the bacterial flagellar motor. Mol Microbiol 78:1117–1129.

21. Zhou J, Lloyd SA, Blair DF. 1998. Electrostatic interactions between rotor and stator in the bacterial flagellar motor. Proc Natl Acad Sci 95:6436–6441.

22. Nord AL, Gachon E, Perez-Carrasco R, Nirody JA, Barducci A, Berry RM, Pedaci F. 2017. Catch bond drives stator mechanosensitivity in the bacterial flagellar motor. Proc Natl Acad Sci 114:12952–12957.

23. Baker AE, O’Toole GA. 2017. Bacteria, rev your engines: Stator dynamics regulate flagellar motility. J Bacteriol 199:e00088–17.

24. Chawla R, Ford KM, Lele PP. 2017. Torque, but not FliL, regulates mechanosensitive flagellar motor-function. Sci Rep 7:5565.

25. Hug I, Deshpande S, Sprecher KS, Pfohl T, Jenal U. 2017. Second messenger-mediated tactile response by a bacterial rotary motor. Science 358:531–534.

26. Lauriano CM, Ghosh C, Correa NE, Klose KE. 2004. The sodium-driven flagellar motor controls exopolysaccharide expression in *Vibrio cholerae*. J Bacteriol 186:4864–4874.

27. Watnick PI, Lauriano CM, Klose KE, Croal L, Kolter R. 2001. The absence of a flagellum leads to altered colony morphology, biofilm development and virulence in *Vibrio cholerae* O139. Mol Microbiol 39:223–235.

28. Wu DC, Zamorano-Sánchez D, Pagliai FA, Park JH, Floyd KA, Lee CK, Kitts G, Rose CB, Bilotta EM, Wong GCL, Yildiz FH. 2020. Reciprocal c-di-GMP signaling: Incomplete flagellum biogenesis triggers c-di-GMP signaling pathways that promote biofilm formation. PLoS Genet 16:e1008703.

29. Hershey DM, Fiebig A, Crosson S. 2021. Flagellar perturbations activate adhesion through two distinct pathways in *Caulobacter crescentus*. MBio 12:10–1128.

30. Harrison JJ, Almblad H, Irie Y, Wolter DJ, Eggleston HC, Randall TE, Kitzman JO, Stackhouse B, Emerson JC, McNamara S, Tyler J. Larsen, Jay Shendure, Lucas R. Hoffman, Daniel J. Wozniak, Matthew R. Parsek. 2020. Elevated exopolysaccharide levels in *Pseudomonas aeruginosa* flagellar mutants have implications for biofilm growth and chronic infections. PLoS Genet 16:e1008848.

31. Lewis KA, Vermilyea DM, Webster SS, Geiger CJ, de Anda J, Wong GCL, O’Toole GA, Hogan DA. 2022. Nonmotile subpopulations of *Pseudomonas aeruginosa* repress flagellar motility in motile cells through a type IV pilus-and Pel-dependent mechanism. J Bacteriol 204:e00528–21.

32. de Anda J, Kuchma SL, Webster SS, Boromand A, Lewis KA, Lee CK, Contreras M, Medeiros Pereira VF, Schmidt W, Hogan DA, O’Hern CS, O’Toole GA, Wong GCL. 2024. How *P. aeruginosa* cells with diverse stator composition collectively swarm. mBio 15:e0332223.

33. Hook AL, Flewellen JL, Dubern J-F, Carabelli AM, Zaid IM, Berry RM, Wildman RD, Russell N, Williams P, Alexander MR. 2019. Simultaneous tracking of *Pseudomonas aeruginosa* motility in liquid and at the solid-liquid interface reveals differential roles for the flagellar stators. mSystems 4:e00390–19.

34. Wu Z, Tian M, Zhang R, Yuan J. 2021. Dynamics of the two stator systems in the flagellar motor of *Pseudomonas aeruginosa* studied by a bead assay. Appl Environ Microbiol 87:e01674–21.

35. Kuchma SL, Delalez NJ, Filkins LM, Snavely EA, Armitage JP, O’Toole GA. 2015. Cyclic di-GMP-mediated repression of swarming motility by *Pseudomonas aeruginosa* PA14 Requires the MotAB stator. J Bacteriol 197:420–430.

36. Baker AE, Webster SS, Diepold A, Kuchma SL, Bordeleau E, Armitage JP, O’Toole GA. 2019. Flagellar stators stimulate c-di-GMP production by *Pseudomonas aeruginosa*. J Bacteriol 201:e00741–18.

37. Baker AE, Diepold A, Kuchma SL, Scott JE, Ha DG, Orazi G, Armitage JP, O’Toole GA. 2016. PilZ domain protein FlgZ mediates cyclic di-GMP-dependent swarming motility control in *Pseudomonas aeruginosa*. J Bacteriol 198:1837–1846.

38. Friedman L, Kolter R. 2004. Genes involved in matrix formation in *Pseudomonas aeruginosa* PA14 biofilms. Mol Microbiol 51:675–690.

39. Ma LZ, Wang D, Liu Y, Zhang Z, Wozniak DJ. 2022. Regulation of biofilm exopolysaccharide biosynthesis and degradation in *Pseudomonas aeruginosa*. Annu Rev Microbiol 76:413–433.

40. Gheorghita AA, Wozniak DJ, Parsek MR, Howell PL. 2023. *Pseudomonas aeruginosa* biofilm exopolysaccharides: assembly, function, and degradation. FEMS Microbiol Rev 47:1–31.

41. Webster SS, Mathelié-Guinlet M, Verissimo AF, Schultz D, Viljoen A, Lee CK, Schmidt WC, Wong GCL, Dufrêne YF, O’Toole GA. 2021. Force-induced changes of PilY1 drive surface sensing by *Pseudomonas aeruginosa*. mBio 13:e0375421.

42. Lee CK, Schmidt WC, Webster SS, Chen JW, O’Toole GA, Wong GCL. 2022. Broadcasting of amplitude-and frequency-modulated c-di-GMP signals facilitates cooperative surface commitment in bacterial lineages. Proc Natl Acad Sci 119:e2112226119.

43. Lee CK, De Anda J, Baker AE, Bennett RR, Luo Y, Lee EY, Keefe JA, Helali JS, Ma J, Zhao K, Golestanian R, O’Toole GA, Wong GCL. 2018. Multigenerational memory and adaptive adhesion in early bacterial biofilm communities. Proc Natl Acad Sci 115:4471–4476.

44. Luo Y, Zhao K, Baker AE, Kuchma SL, Coggan KA, Wolfgang MC, Wong GCL, O’Toole GA. 2015. A hierarchical cascade of second messengers regulates *Pseudomonas aeruginosa* Surface Behaviors. mBio 6:e02456–14.

45. Webster SS, Lee CK, Schmidt WC, Wong GCL, O’Toole GA. 2021. Interaction between the type 4 pili machinery and a diguanylate cyclase fine-tune c-di-GMP levels during early biofilm formation. Proc Natl Acad Sci 118:e2105566118.

46. Lewis KA, Baker AE, Chen AI, Harty CE, Kuchma SL, O’Toole GA, Hogan DA. 2019. Ethanol decreases *Pseudomonas aeruginosa* flagellar motility through the regulation of flagellar stators. J Bacteriol 201:e00285–19.

47. Morimoto YV, Nakamura S, Hiraoka KD, Namba K, Minamino T. 2013. Distinct roles of highly conserved charged residues at the MotA-FliG interface in bacterial flagellar motor rotation. J Bacteriol 195:474–481.

48. Zhou J, Sharp LL, Tang HL, Lloyd SA, Billings S, Braun TF, Blair DF. 1998. Function of protonatable residues in the flagellar motor of *Escherichia coli*: a critical role for Asp 32 of MotB. J Bacteriol 180:2729–2735.

49. Nakamura S, Minamino T, Kami-Ike N, Kudo S, Namba K. 2014. Effect of the MotB(D33N) mutation on stator assembly and rotation of the proton-driven bacterial flagellar motor. Biophysics 10:35–41.

50. Kuchma SL, Ballok AE, Merritt JH, Hammond JH, Lu W, Rabinowitz JD, O’Toole GA. 2010. Cyclic-di-GMP-mediated repression of swarming motility by *Pseudomonas aeruginosa*: The *pilY1* gene and its impact on surface-associated behaviors. J Bacteriol 192:2950–64.

51. Merritt JH, Ha D-G, Cowles KN, Lu W, Morales DK, Rabinowitz J, Gitai Z, O’Toole GA. 2010. Specific control of *Pseudomonas aeruginosa* surface-associated behaviors by two c-di-GMP diguanylate cyclases. mBio 1:e00183–10.

52. Merritt JH, Brothers KM, Kuchma SL, O’Toole GA. 2007. SadC reciprocally influences biofilm formation and swarming motility via modulation of exopolysaccharide production and flagellar function. J Bacteriol 189:8154–64.

53. Guan C, Huang Y, Zhou Y, Han Y, Liu S, Liu S, Kong W, Wang T, Zhang Y. 2024. FlhF affects the subcellular clustering of WspR through HsbR in *Pseudomonas aeruginosa*. Appl Environ Microbiol 90:e0154823.

54. Pandza S, Baetens M, Park CH, Au T, Keyhan M, Matin A. 2000. The G-protein FlhF has a role in polar flagellar placement and general stress response induction in *Pseudomonas putida*. Mol Microbiol 36:414–423.

55. Murray TS, Kazmierczak BI. 2006. FlhF is required for swimming and swarming in *Pseudomonas aeruginosa*. J Bacteriol 188:6995–7004.

56. Dasgupta N, Wolfgang MC, Goodman AL, Arora SK, Jyot J, Lory S, Ramphal R. 2003. A four-tiered transcriptional regulatory circuit controls flagellar biogenesis in *Pseudomonas aeruginosa*. Mol Microbiol 50:809–824.

57. Zhang L, Wu Z, Zhang R, Yuan J. 2022. FliL differentially interacts with two stator systems to regulate flagellar motor output in *Pseudomonas aeruginosa*. Appl Environ Microbiol 88:e01539–22.

58. Nakamura S, Minamino T. 2019. Flagella-driven motility of bacteria. Biomolecules 9:279.

59. Kuchma SL, Griffin EF, O’Toole GA. 2012. Minor pilins of the type IV pilus system participate in the negative regulation of swarming motility. J Bacteriol 194:5388–403.

60. Nicastro GG, Kaihami GH, Pulschen AA, Hernandez-Montelongo J, Boechat AL, de Oliveira Pereira T, Rosa CGT, Stefanello E, Colepicolo P, Bordi C, Baldini RL. 2020. c-di-GMP-related phenotypes are modulated by the interaction between a diguanylate cyclase and a polar hub protein. Sci Rep 10:3077–3077.

61. Semmler ABT, Whitchurch CB, Leech AJ, Mattick JS. 2000. Identification of a novel gene, fimV, involved in twitching motility in Pseudomonas aeruginosa. Microbiology 146:1321– 1332.

62. Inclan YF, Persat A, Greninger A, Von Dollen J, Johnson J, Krogan N, Gitai Z, Engel JN. 2016. A scaffold protein connects type IV pili with the Chp chemosensory system to mediate activation of virulence signaling in *Pseudomonas aeruginosa*. Mol Microbiol 101:590–605.

63. Wehbi H, Portillo E, Harvey H, Shimkoff AE, Scheurwater EM, Howell PL, Burrows LL. 2011. The peptidoglycan-binding protein FimV promotes assembly of the *Pseudomonas aeruginosa* type IV pilus secretin. J Bacteriol 193:540–50.

64. Laventie B-J, Sangermani M, Estermann F, Manfredi P, Planes R, Hug I, Jaeger T, Meunier E, Broz P, Jenal U. 2019. A surface-induced asymmetric program promotes tissue colonization by *Pseudomonas aeruginosa*. Cell Host Microbe 25:140–152.

65. Kuchma SL, Brothers KM, Merritt JH, Liberati NT, Ausubel FM, O’Toole GA. 2007. BifA, a cyclic-di-GMP phosphodiesterase, inversely regulates biofilm formation and swarming motility by *Pseudomonas aeruginosa* PA14. J Bacteriol 189:8165–78.

66. Caiazza NC, O’Toole GA. 2004. SadB is required for the transition from reversible to irreversible attachment during biofilm formation by *Pseudomonas aeruginosa* PA14. J Bacteriol 186:4476–4485.

67. Caiazza NC, Merritt JH, Brothers KM, O’Toole GA. 2007. Inverse regulation of biofilm formation and swarming motility by *Pseudomonas aeruginosa* PA14. J Bacteriol 189:3603– 3612.

68. Persat A, Inclan YF, Engel JN, Stone HA, Gitai Z. 2015. Type IV pili mechanochemically regulate virulence factors in *Pseudomonas aeruginosa*. Proc Natl Acad Sci 112:7563– 7568.

69. Geiger CJ, O’Toole GA. 2023. Evidence for the Type IV pilus retraction motor PilT as a component of the surface sensing system in *Pseudomonas aeruginosa*. J Bacteriol 205:e0017923.

70. Manner C, Dias Teixeira R, Saha D, Kaczmarczyk A, Zemp R, Wyss F, Jaeger T, Laventie B-J, Boyer S, Malone JG, Qvortrup K, Andersen JB, Givskov M, Tolker-Nielsen T, Hiller S, Drescher K, Jenal U. 2023. A genetic switch controls *Pseudomonas aeruginosa* surface colonization. Nat Microbiol 8:1520–1533.

71. Van Loon JC, Whitfield GB, Wong N, O’Neal L, Henrickson A, Demeler B, O’Toole GA, Parsek MR, Howell PL. 2024. Binding of GTP to BifA is required for the production of Pel-dependent biofilms in *Pseudomonas aeruginosa*. J Bacteriol 206:e0033123.

72. Dahlstrom KM, Giglio KM, Collins AJ, Sondermann H, O’Toole GA. 2015. Contribution of physical interactions to signaling specificity between a diguanylate cyclase and its effector. mBio 6:e01978–15.

73. Köhler T, Curty LK, Barja F, van Delden C, Pechère JC. 2000. Swarming of *Pseudomonas aeruginosa* is dependent on cell-to-cell signaling and requires flagella and pili. J Bacteriol 182:5990–5996.

74. Gibson DG, Young L, Chuang R-Y, Venter JC, Hutchison CA 3rd, Smith HO. 2009. Enzymatic assembly of DNA molecules up to several hundred kilobases. Nat Methods 6:343–345.

75. Shanks RMQ, Caiazza NC, Hinsa SM, Toutain CM, O’Toole GA. 2006. *Saccharomyces cerevisiae*-based molecular tool kit for manipulation of genes from gram-negative bacteria. Appl Environ Microbiol 72:5027–5036.

76. Schweizer HP. 1992. Allelic exchange in *Pseudomonas aeruginosa* using novel ColE1-type vectors and a family of cassettes containing a portable oriT and the counter-selectable *Bacillus subtilis sacB* marker. Mol Microbiol 6:1195–1204.

77. Kulasekara HD, Ventre I, Kulasekara BR, Lazdunski A, Filloux A, Lory S. 2005. A novel two-component system controls the expression of *Pseudomonas aeruginosa* fimbrial cup genes. Mol Microbiol 55:368–380.

78. Ha DG, Kuchma SL, O’Toole GA. 2014. Plate-based assay for swarming motility in *Pseudomonas aeruginosa*. Methods Mol Biol. 1149:67–72.

